# Pyroptosis-related gene signatures can robustly diagnose skin cutaneous melanoma and predict the prognosis

**DOI:** 10.1101/2021.04.17.440259

**Authors:** Anji Ju, Jiaze Tang, Shuohua Chen, Yan Fu, Yongzhang Luo

## Abstract

Skin cutaneous melanoma (SKCM) is a chronically malignant tumor with a high mortality rate. Pyroptosis, a kind of pro-inflammatory programmed cell death, has been linked to cancer in recent studies. However, the value of pyroptosis in the diagnosis and prognosis of SKCM is not clear. In this study, it was discovered that 20 pyroptosis-related genes (PRGs) differed in expression between SKCM and normal tissues, which were related to diagnosis and prognosis. Firstly, based on these genes, nine machine-learning algorithms were shown to perform well in constructing diagnostic classifiers, including K-Nearest Neighbor (KNN), logistic regression, Support Vector Machine (SVM), Artificial Neural Network (ANN), decision tree, random forest, XGBoost, LightGBM, and CatBoost. Secondly, the least absolute shrinkage and selection operator (LASSO) Cox regression analysis was applied and the prognostic model was constructed based on 9 PRGs. Subgroups in low and high risks determined by the prognostic model were shown to have different survival. Thirdly, functional enrichment analyses were performed by applying the gene set enrichment analysis (GSEA), and results suggested that the risk was related to immune response. Finally, immune infiltration analysis was carried out, which showed that fractions of activated CD4+ memory T cells, γδ T cells, M1 macrophages, and M2 macrophages were significantly different between subgroups. In conclusion, the expression signatures of pyroptosis-related genes are effective and robust in the diagnosis and prognosis of SKCM, which is related to immunity.

## 1 Introduction

Malignant skin cutaneous melanoma (SKCM) is a serious life-threatening disease, and the incidence rate of SKCM is rapidly increasing throughout the world (1, 2). SKCM lacks specific treatment other than early surgical resection, which leads to a poor prognosis and extremely high mortality (3). Although non-Caucasian populations are less likely to develop melanoma, the severity of SKCM in Africa, Asia, Central America, and South America has increased (4). Lack of prevention and early diagnosis programs may contribute to the increased prevalence of SKCM in these regions (5). Therefore, developing efficient diagnosis and prognosis methods is important for the treatment of SKCM.

Pyroptosis, or caspase 1-dependent cell death, also known as cellular inflammatory necrosis, is triggered by various pathological stimuli, such as microbial infections, stroke, heart attack, and cancer (6). The term *pyroptosis* was first proposed in 2001 from the Greek roots *pyro*, relating to fire or fever, and *ptosis (to-sis)* to denote a falling, to describe pro-inflammatory programmed cell death (7). In addition to apoptosis, ferroptosis, and autophagy, this new type of cell death has become a hot spot recently.

Pyroptosis is characterized by the rapid rupture of the plasma membrane and the release of pro-inflammatory intracellular contents. A canonical pathway of pyroptosis is triggered by the activation of inflammasomes which are cytoplasmic multi-protein platforms containing the nucleotide-binding oligomerization domain (NOD)-like receptor (NLR) family (8). Caspase-1 can be activated by inflammasomes, which leads to the cleavage of gasdermin D (GSDMD) and both the maturation and secretion of pro-inflammatory cytokines, such as IL-18 and IL-1B (9). Caspase-1-dependent plasma membrane pores dissipate cellular ion gradients, resulting in osmotic pressure increase, which leads to water influx and cell swelling (10). Ultimately, osmotic lysis occurs and inflammatory intracellular contents are released (10). Caspase-1 dependence is a defining feature of pyroptosis in which mediates cell lysis during pyroptosis and is not involved in apoptosis (11–13). Besides GSDMD, the plasma membrane pores formation can be executed by the cleavage of other gasdermin proteins, especially gasdermin E (GSDME) which can be cleaved by caspase-3 to trigger pyroptosis (14, 15).

The mechanism and functions of pyroptosis in tumor cells have been extensively studied, but its relationship to cancer prognosis has been ambiguous. This is because pyroptosis plays a dual role in cancer progression. On one hand, inducing pyroptosis may be a feasible method to kill tumor cells; on the other hand, as a type of pro-inflammatory death, pyroptosis can form a suitable microenvironment for tumor cell growth and thus promote tumor growth (16–20). Studying the relationship between pyroptosis and clinical features of SKCM is helpful for its treatment, but the value of pyroptosis in the diagnosis and prognosis of SKCM has not been reported. Therefore, in this systematic study, classifiers were built through machine-learning algorithms to mine out the diagnosis value of pyroptosis-related genes (PRGs) in distinguishing between SKCM and normal tissue. Then a novel PRGs prognostic risk signature in SKCM was constructed for survival predicting. Besides, prognostic risk-related phenotypes including immune infiltration were analyzed. Thus, this study provides both a novel understanding of the role of pyroptosis and novel methods for the diagnosis and prognosis prediction in SKCM.

## 2 Materials and methods

### 2.1 Data collection

The study design and grouping are shown in Figure 1. Transcriptome profiles and clinical data in SKCM patients were collected in the database of The Cancer Genome Atlas (TCGA) -SKCM (18^th^ December 2019. https://portal.gdc.cancer.gov/) and Gene Expression Omnibus (GEO. https://www.ncbi.nlm.nih.gov/geo/) including GSE54467, GSE65904, GSE98394, and GSE112509 (21–25). Transcriptome profiles in normal skin tissues were collected in the database of Genotype-Tissue Expression Project (GTEx-SKIN. https://gtexportal.org/home/). The RNA-seq data in TCGA-SKCM, GSE98394, GSE112509, and GTEx-SKIN were converted to Transcripts per Kilobase Million (TPM) format. The microarray data in GSE54467 and GSE65904 were normalized by using the R package “limma”. Repeat values were averaged and missing values were removed. The RNA-seq data in TCGA-SKCM and GTEx-SKIN were merged and normalized by using the R package “limma”.

**Figure 1:**
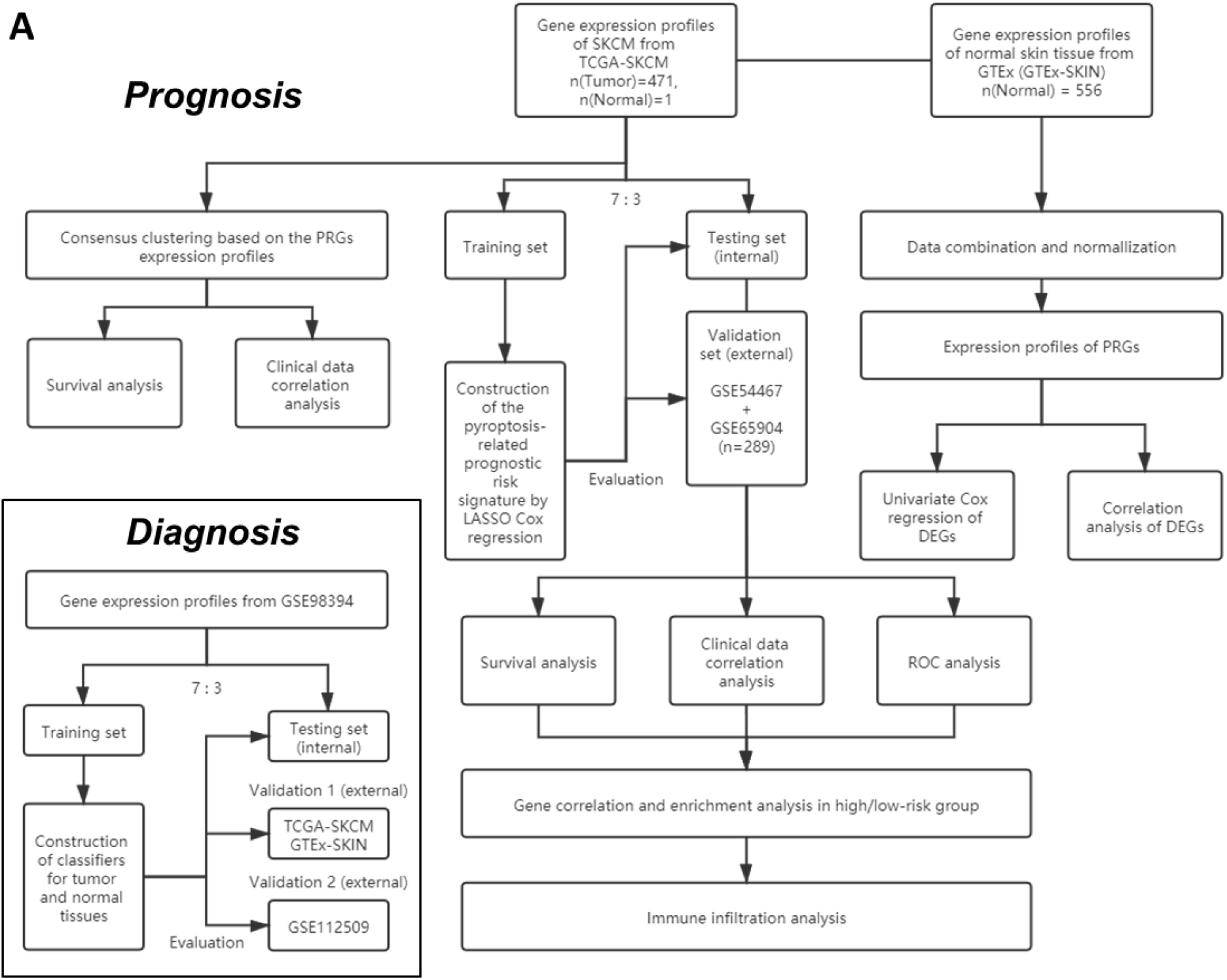
The flowchart of the overall procedures. This flowchart illustrates the process of data collection and analyses for diagnostic and prognostic studies.

### 2.2 Identification of differentially expressed genes

Twenty PRGs (listed in Table S1) were retrieved in the GeneCards database (8th January 2020. https://www.genecards.org/) by the keyword “pyroptosis” and verified in several reviews (26–29). The “limma” package was used to identify differentially expressed genes (DEGs) between SKCM and normal tissues with the *p*-value <0.05. The correlation of DEGs was analyzed and demonstrated by using the R package “corrplot”. The significance of relationships between OS and the DEGs in TCGA-SKCM was determined using univariate Cox regression analysis, which was carried out by using the “survival” R package. A protein-protein interaction (PPI) network for the DEGs was obtained from Search Tool for the Retrieval of Interacting Genes (STRING v11.0, https://string-db.org/).

### 2.3 Construction and evaluation of PRGs-based classifiers for SKCM diagnosis

Data from GSE98394 were randomly divided into a training set and a testing set according to 7:3. Data from the training set were used to train the classifiers respectively based on the K-Nearest Neighbor (KNN), logistic regression, Support Vector Machine (SVM), Artificial Neural Network (ANN), decision tree, random forest, XGBoost, LightGBM, and CatBoost *via* following Python packages: Scikit-learn (sklearn) v0.23.2, XGBoost v1.3.3, LightGBM v3.1.1, and CatBoost v0.24.4 (30–33).

The “sklearn.metrics” Python package was used to evaluate the PRGs-based classifiers, and the “matplotlib” Python package was used to plot the receiver operating characteristic (ROC) curves. Besides the area under ROC curves (AUC), accuracy, precision (also known as positive predictive value), recall (also known as sensitivity), and F1 score were calculated to evaluate the prediction performance of the models by using the “sklearn.metrics” Python package. To assess the quality of the models, the Gini index and Kolmogorov–Smirnov (KS) value were calculated according to the methods described previously (34).

Data from the testing set were used to perform internal evaluations and parameter tuning. Major parameters used in the above algorithms are listed in Table S2. For external evaluations, data from TCGA-SKCM & GTEx-SKIN (validation 1 set) and GSE112509 (validation 2 set) were used. Data from each group were normalized by employing the “StandardScaler” function from the “sklearn.preprocessing” Python package before training and evaluations.

### 2.4 Consensus clustering analysis of PRGs

To classify the SKCM by consensus clustering, R packages “limma” and “ConsensusClusterPlus” were used. The “prcomp” function in the “stats” R package was used to conduct principal component analysis (PCA) based on the clusters. The correlations between clusters and clinical characteristics, including overall survival (OS), were analyzed by employing the chi-square test and R package “survival”. The results were presented by heat maps and Kaplan-Meier (KM) curves via R packages “pheatmap” and “survival”.

### 2.5 Construction of PRGs-based SKCM prognostic model

The least absolute shrinkage and selection operator (LASSO) Cox regression analysis was performed by using the R package “glmnet” to narrow down the candidate genes and to develop the prognostic model. The penalty parameter (λ) was determined by the minimum parameters. The risk scores were calculated using the following equations:

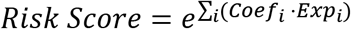

where *Coef* is the coefficient and *Exp* is the expression level of every retained gene. Data from TCGA-SKCM were randomly divided into a training set and a testing set according to 7:3. The risk score was calculated by using the data from the training set. Data from the testing set was used for the internal evaluation. Data from GSE54467 and GSE65904 were merged and normalized as a validation set by using the R package “limma” for the external evaluation. The R package “survivalROC” was employed to perform 3- and 5-year ROC analysis.

The correlation between subgroups and clinical characteristics in TCGA-SKCM was analyzed by employing the chi-square test and presented by heat map. The independence of the clinical factors and the risk score calculated from the prognostic model was determined using univariate and multivariate Cox regression analyses, which were carried out by using the “survival” R package.

### 2.6 Gene sets enrichment analysis

The DEGs (|log2FC| ≥ 1 and FDR < 0.05) between the low- and high-risk subgroups in TCGA-SKCM were filtered, which was carried out with the Gene Ontology (GO) analysis by using the “clusterProfiler” R package. Besides, gene set enrichment analysis (GSEA) was used in TCGA-SKCM to identify the biological processes that were significantly alerted between the high-risk and low-risk subgroups (35, 36). The Java GSEA software (version 4.0.1) was employed and the gene set “c2.cp.kegg.v7.4.symbols.gmt” from the database of Kyoto Encyclopedia of Genes and Genomes (KEGG) was chosen as the reference (37–39). Biological processes with the normalized *p* < 0.05 and the false discovery rate (FDR) q value < 0.05 were considered as statistically significant. The top biological processes that had been altered were chosen based on a ranking of normalized enrichment ratings (NES).

### 2.7 Immune infiltration analysis

Transcriptome data from TCGA-SKCM was transformed into the total abundance of immune cells by utilizing the Cell-type Identification by Estimating Relative Subsets of RNA Transcripts (CIBERSORT) analysis with the “CIBERSORT” R package (40, 41). Patients were divided into low- and high-infiltration subgroups according to the median level. Tumor IMmune Estimation Resource 2.0 (TIMER2.0, http://timer.cistrome.org/) was employed to analyze the correlation between the immune infiltration and OS in SKCM (42–44).

### 2.8 Statistical analyses

Wilcoxon test was applied to compare the gene expression levels between the normal skin and SKCM tissues and the immune infiltration levels between subgroups. The two-sided log-rank test was used to compare the OS between subgroups. Other statistical methods are specifically described above. All statistical analyses were accomplished with R (v3.6.2) and Anaconda 3 (Python v3.8.5).

## 3 Results

### 3.1 Identification of differentially expressed PRGs between normal skin and SKCM tissues

Expression levels of 20 PRGs were compared between 557 normal and 471 tumor tissues from GTEx-SKIN and TCGA-SKCM data. It was observed that all the 20 PRGs were significantly differentially expressed (all *p* < 0.01. **Figure 2A** and **Figure S1A**). Among them, 11 genes (CASP1, PYCARD, APIP, FOXO3, IL18, GSDMA, GSDMC, CASP4, GSDMB, NLRP1, and NAIP) were downregulated while 9 genes (NLRP9, DHX9, CASP3, NLRC4, AIM2, NLRP3, IL1B, GSDME, and GSDMD) were upregulated in tumor tissues. In addition, 13 genes showed significant associations with OS (**Figure 2B**). Among them, 11 genes were protecting factors (hazard ratio < 1) and 2 genes were risk factors (hazard ratio > 1).

**Figure 2:**
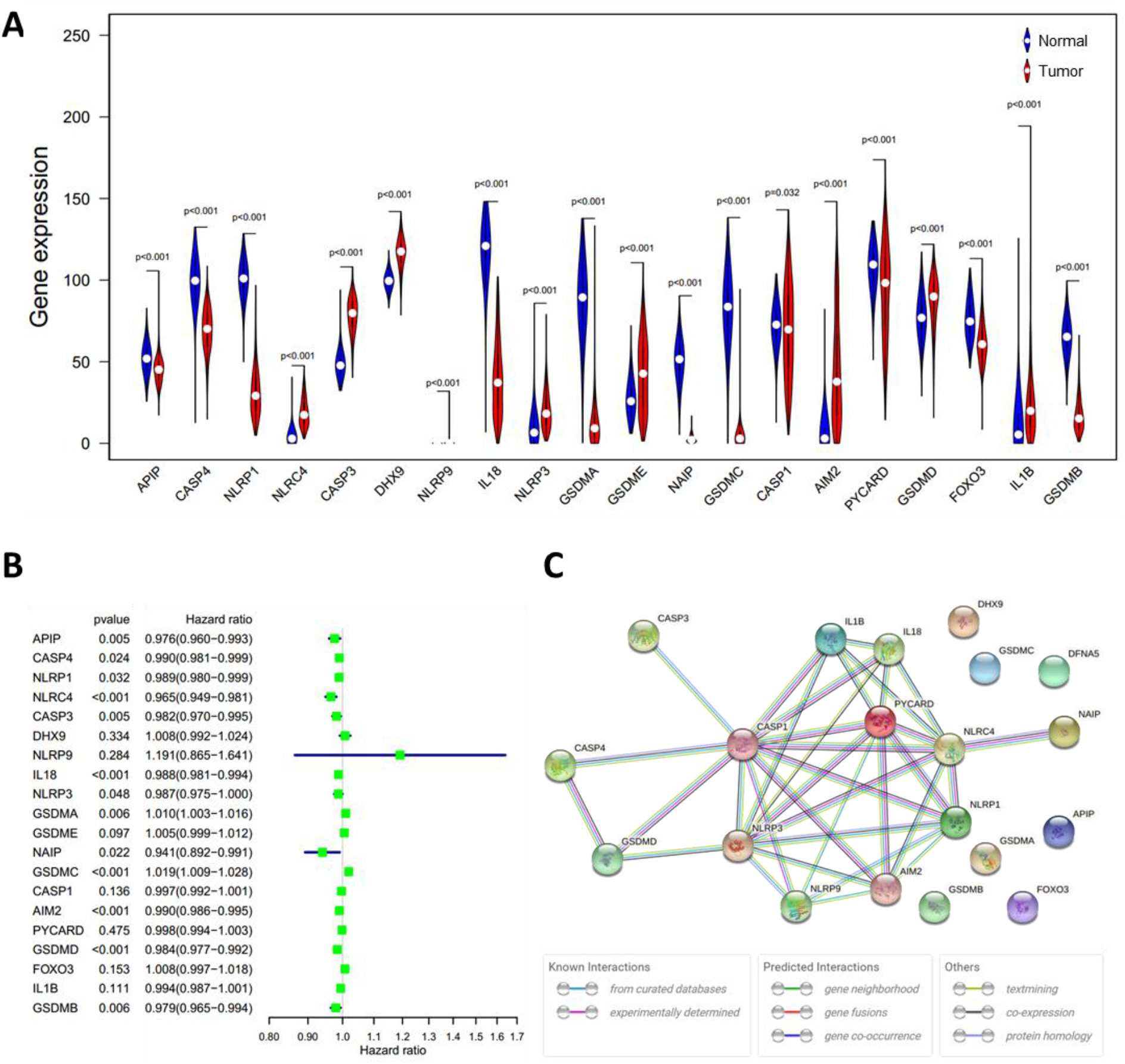
Expressions and the associations with OS of the 20 PRGs. **(A)** Violin plot of PRGs between the normal (blue) and the tumor tissues (red). **(B)** Significance and hazard ratio values of OS-related PRGs in univariate Cox regression. **(C)** PPI network showing the interactions of the PRGs (interaction score = 0.9). The bottom boxes show the types of interactions.

To further explore the interactions of these PRGs, PPI and expression correlation analysis were performed (**Figure 2C** and **Figure S1B**). The minimum required interaction score for the PPI analysis was set at 0.9 (the highest confidence). The results suggested that CASP1, CASP3, GSDMD, NLRP3, PYCARD, AIM2, and NLRC4 play central roles in the pyroptosis process of SKCM. The Human Protein Atlas (www.proteinatlas.org) was used to retrieve immunohistochemistry staining images of proteins encoded by PRGs in SKCM, showing cellular sublocalization of these molecules (**Figure S2**). It can be seen that several widely reported pyroptosis-related proteins were in high levels, including AIM2, CASP1, CASP3, GSDMD, and GSDME, which indicate that pyroptosis occurred in a large part of SKCM tissues.

### 3.2 Diagnosis value of PRGs-based classifiers in SKCM

Given the significant difference in PRG expression between normal and tumor tissues, it was hypothesized that PRGs can be used to diagnose SKCM. To verify this hypothesis, nine commonly used machine-learning algorithms were used to construct diagnostic classifiers, including KNN, logistic regression, SVM, ANN, decision tree, random forest, XGBoost, LightGBM, and CatBoost. Data from GSE98394 were randomly divided into a training set and a testing set according to 7:3. Respectively, classifiers based on the above algorithms were trained by using the RNA-seq data in the training set. The testing set was designed to perform internal evaluations. As expected, RNA-seq data of PRGs were suitable for building the SKCM diagnostic classifiers, because of the high accuracy in the training and testing set (**Table 1, Table S4,** and **Figure 3J**).

**Figure 3:**
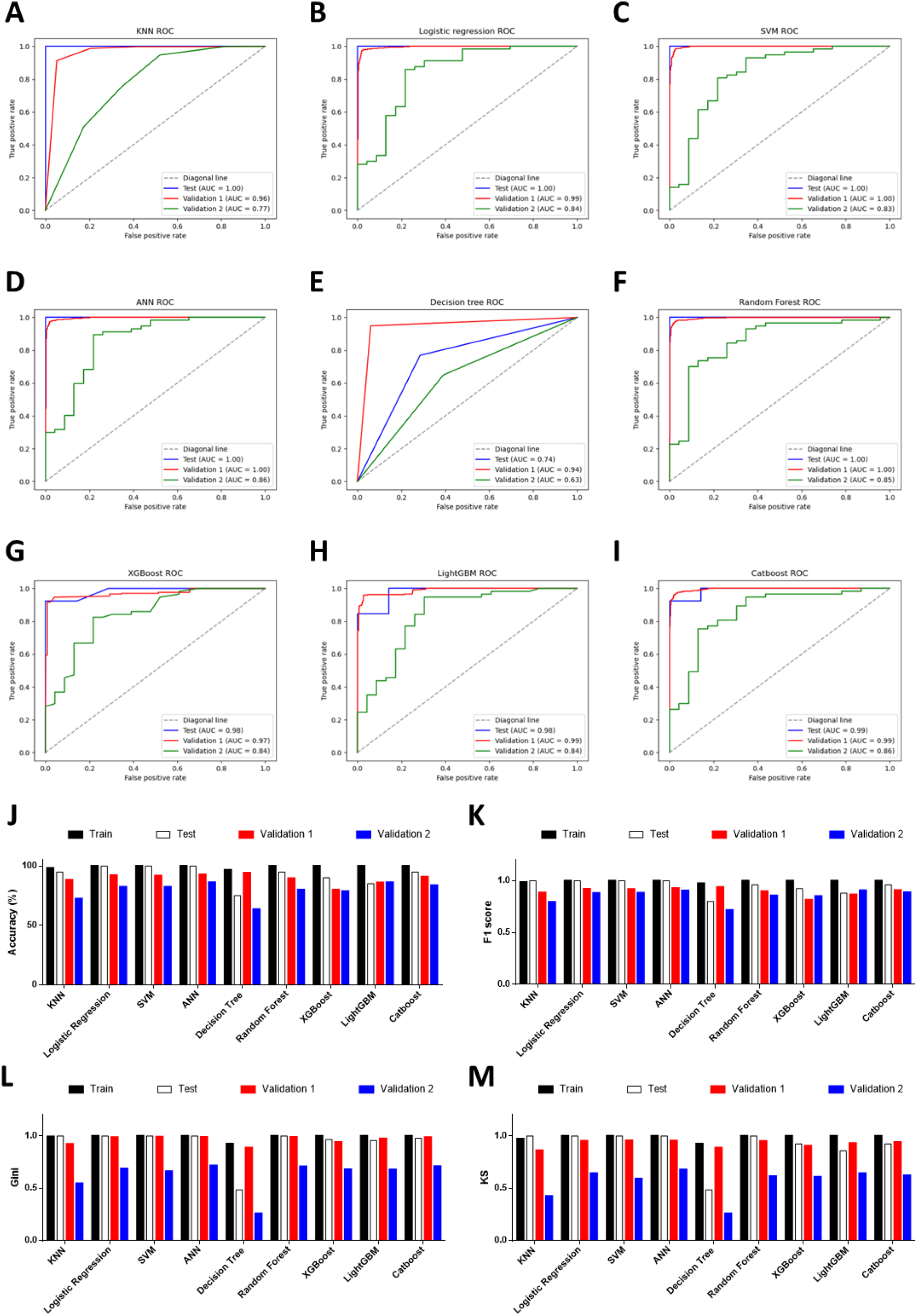
Performance evaluations of diagnostic classifiers based on 9 algorithms. **(A-I)** ROC curves for evaluating the predictive performance of the diagnostic models respectively based on K-Nearest Neighbor (A), logistic regression (B), Support Vector Machine (C), Artificial Neural Network (D), decision tree (E), random forest (F), XGBoost (G), LightGBM (H), and CatBoost (I). Data from GSE98394 were randomly divided into a training set (not shown due to the AUCs were extremely close to 1.0 in all classifiers) and a testing set (blue line) according to 7:3. Validation 1: the combination of TCGA-SKCM & GTEx-SKIN (red line). Validation 2: GSE112509 (green line). **(J)** Columns showing the accuracy (%) of each classifier in different datasets. **(K)** Columns showing the F1 score of each classifier in different datasets. **(L)** Columns showing the Gini index of each classifier in different datasets. **(M)** Columns showing the KS value of each classifier in different datasets.

**Table 1:**
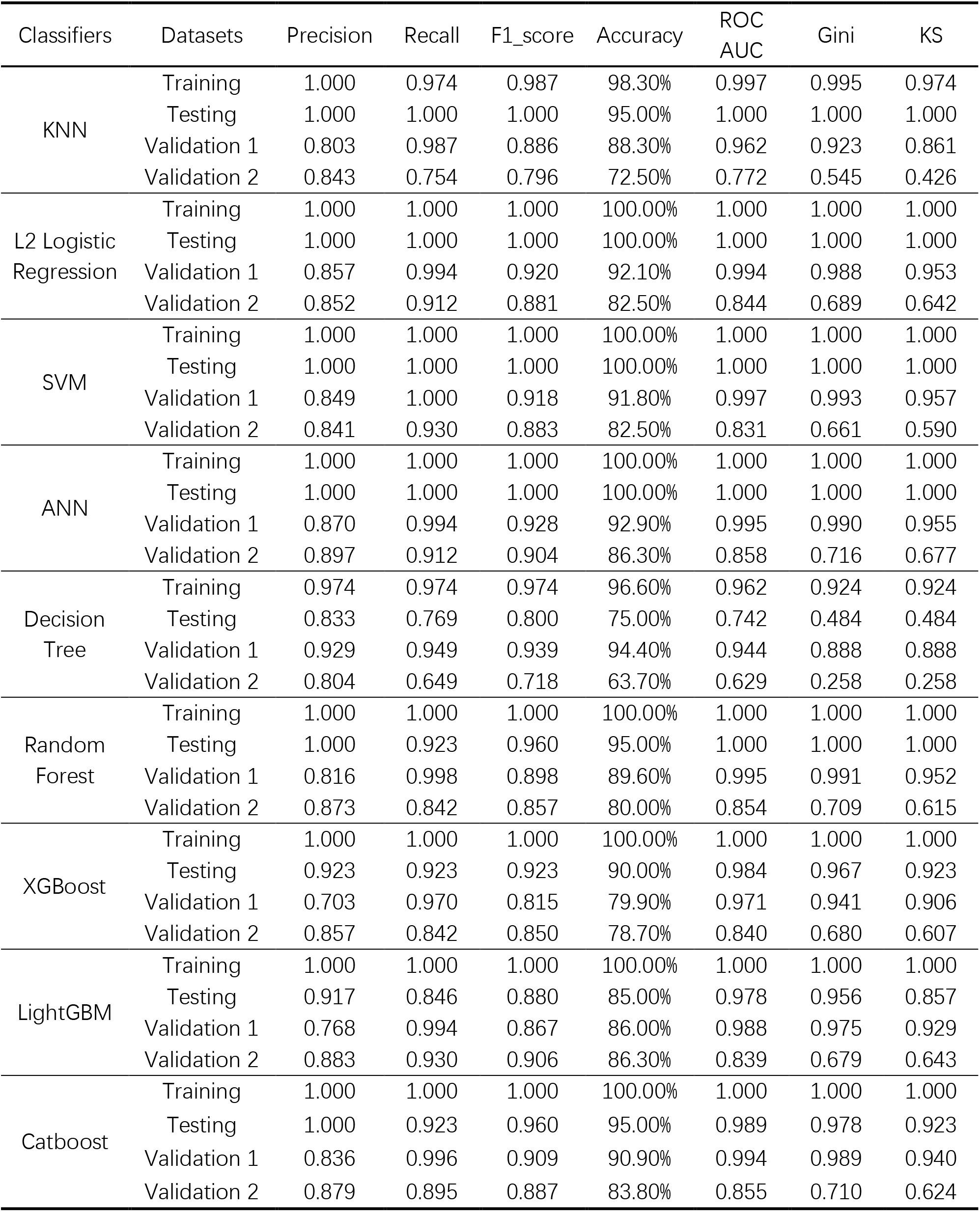
Evaluation parameters of classifiers in different datasets.

In addition to the accuracy, ROC curves were used to evaluate the sensitivity and specificity of the classifiers. In the testing set, except for the poor performance of the decision tree, the AUC values of the other eight algorithms were all higher than 0.900 (**Table 1**). This suggests that PRGs had a very high capacity to distinguish between normal and tumor samples in a single study (GSE98394). To verify the performance of these classifiers in out-of-sample data with different sample sizes and levels of balance, two external validation sets were used to perform ROC analysis. In the validation set 1 (TCGA-SKCM & GTEx-SKIN) with a relatively larger and more balanced sample size (the numbers of tumor and normal samples are 471 and 557), all classifiers performed well (**Figure 3A-I,** red line). Furthermore, except for KNN and decision tree, classifiers based on the other seven algorithms worked well in the validation set 2 (GSE112509) with a relatively smaller and unbalanced sample size (the numbers of tumor and normal samples are 57 and 23) (**Figure 3A-I,** green line).

In order to further evaluate the classifiers, precision, recall and F1 score were calculated and the results were consistent with the ROC analysis (**Table 1** and **Figure 3K**). Moreover, the Gini index and KS value were estimated to confirm the results (**Table 1,** and **Figure 3L, M**). Commonly, when these parameters are close to 1.000, it indicates that the classifier has a strong ability to distinguish. Besides, the classifier is robust when the difference of these parameters among datasets is minimal. Considering all the evaluation parameters, it was found that ANN is the most suitable algorithm to construct the diagnostic model based on PRGs in this study, while logistic regression, random forest, and SVM also performed well, which suggests that the expression signature of PRGs has a high diagnostic benefit in SKCM.

### 3.3 Identification of SKCM clusters using consensus clustering

In order to investigate the therapeutic utility of PRGs, we attempted to divide the SKCM samples into clusters depending on gene expression patterns (**Figure S3**). The number of clusters was represented by the letter “k”. The empirical CDF was plotted to determine the optimum k value for the sample distribution to reach maximal stability (**Figure S3A, B**). Consensus matrices showed that, with k = 2, patients in TCGA-SKCM could be divided into two distinct and non-overlapping clusters, which was verified by the PCA (**Figure S3C** and **Figure 4A**). It was observed that there are significant differences in OS and the stage of SKCM (**Figure 4B, C**). As shown in Figure 4B, cluster 1 (n = 189) had a significantly better OS than cluster 2 (n = 282).

**Figure 4:**
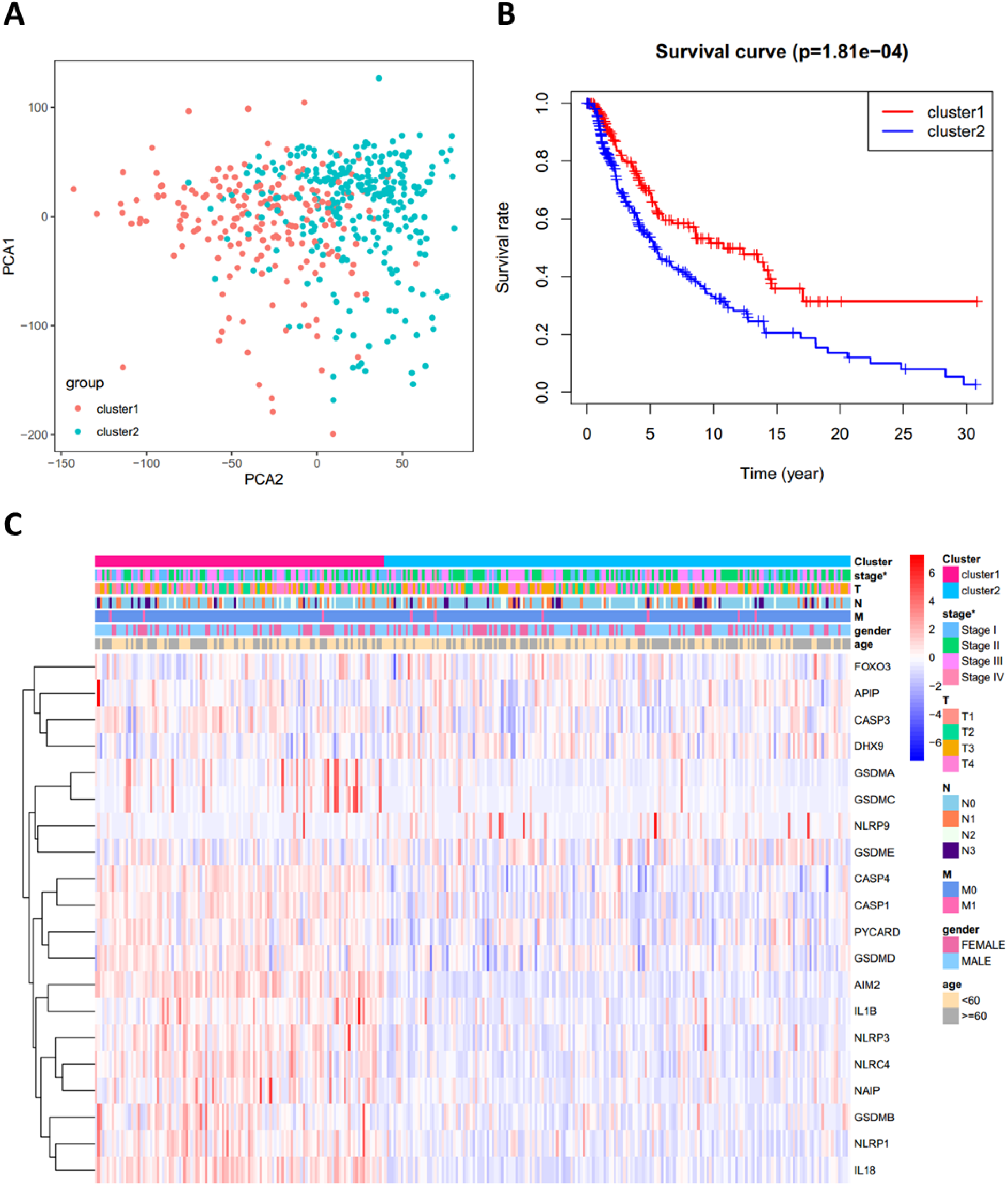
Consensus clustering analysis of PRGs. **(A)** PCA plot for clusters. **(B)** KM curves showing the OS of the two clusters **(C)** Heatmap and the clinical characters of the two clusters (T, N, and M are the tumor node metastasis classification) (* *p* < 0.05).

### 3.4 Prognostic value of PRGs expression signature in SKCM

Cox regression analysis was used to evaluate the correlations between each PRG and survival status to assess the prognostic value of PRGs expression signature. Data from TCGA-SKCM were randomly divided into a training set and a testing set according to 7:3. To narrow down the candidate genes and construct the prognostic model, the LASSO Cox regression model was used in the training set. Nine genes and their coefficients (**Table 2**) were eventually preserved, and the penalty parameter (λ) was determined by the minimum parameters (**Figure 5A, B**). Data from GSE54467 and GSE65904 were merged and normalized as a validation set for the external evaluation. The risk scores in the test and validation sets were calculated by the same equation obtained from the training set (**Figure S4**).

**Figure 5:**
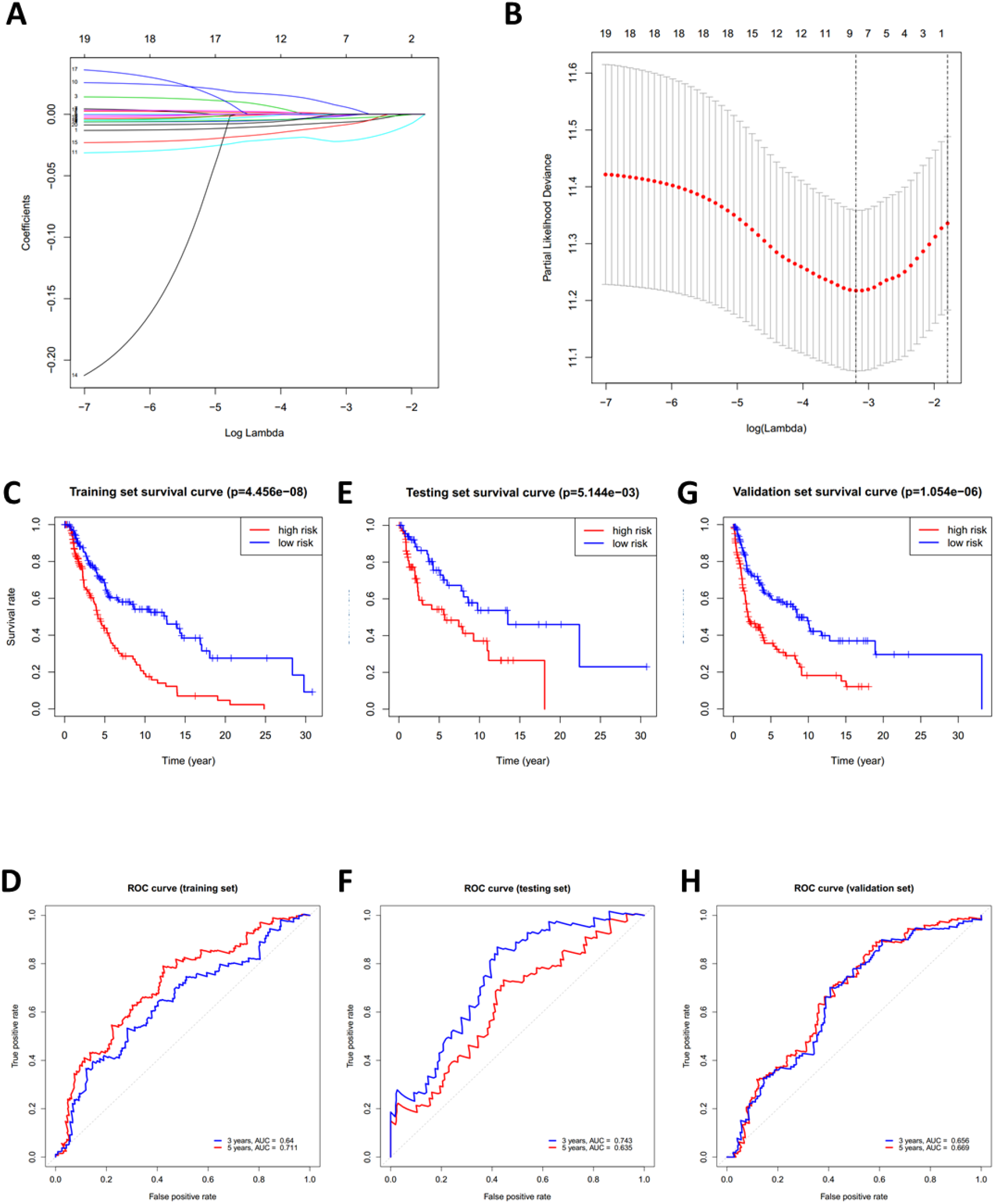
Construction of the PRGs-based prognostic model. **(A)** LASSO regression of the 7 OS-related genes. **(B)** Cross-validation for tuning the parameter in the LASSO regression. **(C-H)** KM curves showing the OS of the low- and high-risk subgroups. ROC curves demonstrated the predictive efficiency of the risk score for 3- and 5-year survival. Data from TCGA-SKCM were randomly divided into a training set (C, D) and a testing set (E, F) according to 7:3. GSE54467 and GSE65904 were merged as the validation set (G, H).

**Table 2:** Coefficients in the LASSO Cox regression model.

| *i* | Gene | Coef |
| --- | --- | --- |
| 1 | GSDMD | -0.006861 |
| 2 | GSDME | 0.0003969 |
| 3 | CASP4 | -0.001943 |
| 4 | GSDMC | 0.0079361 |
| 5 | NLRC4 | -0.022123 |
| 6 | APIP | -0.009636 |
| 7 | AIM2 | -0.003569 |
| 8 | CASP3 | -0.00106 |
| 9 | IL18 | -0.000169 |

According to the median risk score, patients in the training set were divided into low- and high-risk subgroups, and a significant difference in OS was observed *via* the KM survival analysis (**Figure 5C**). The lifespan of patients in the high-risk subgroup was shorter than those in the low-risk subgroup. The sensitivity and specificity of the prognostic model were determined using the time-dependent ROC analysis, and the AUC was 0.640 for 3-year survival and 0.711 for 5-year survival, respectively (**Figure 5D**). Furthermore, patients in the test and validation sets were also divided according to the median risk score. The OS and ROC analyses of these two subgroups showed similar results to the training set (**Figure 5E-H**).

In addition, significant differences in the tumor stage were observed between low-and high-risk subgroups, such as more stage-IV and fewer T1 samples in the high-risk subgroup (**Figure 6A**). This observation led us to wonder whether the risk score could function as an independent prognostic factor in SKCM. To prove this hypothesis, univariate and multivariable Cox regression analyses were performed, which indicated that the PRGs-based prognostic risk was indeed an independent factor in predicting poor survival in TCGA-SKCM (**Figure 6B, C**). These results suggest that the PRGs-based prognostic model is robust and independent in predicting the prognosis of SKCM.

**Figure 6:**
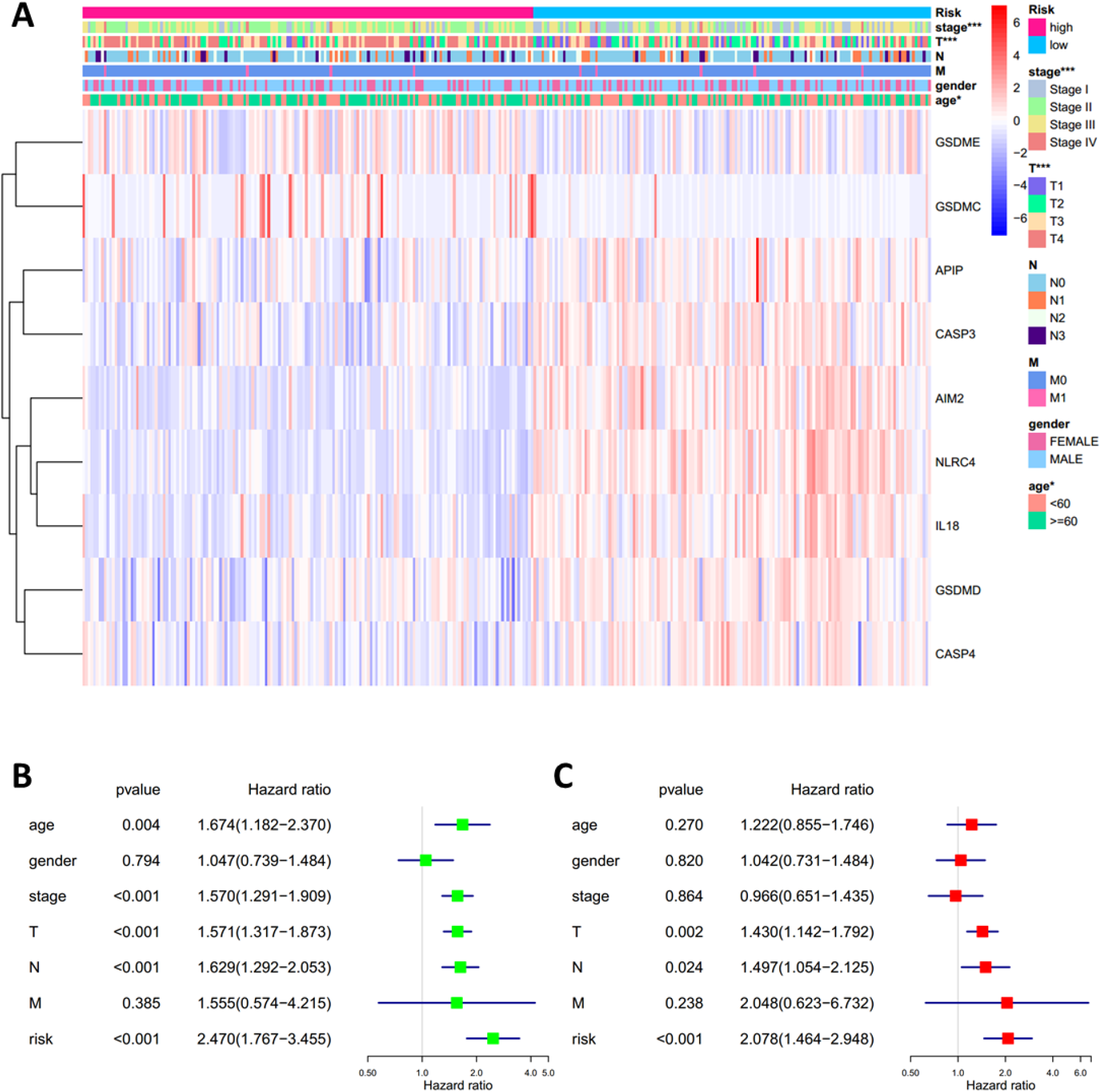
Univariate and multivariate Cox regression analyses for the risk score. **(A)** Heatmap and the clinical characters of low- and high-risk subgroups (T, N, and M are the tumor node metastasis classification) (\**p* < 0.05, \*\**p* < 0.01, *** *p* < 0.001). **(B-C)** Univariate (B) and multivariate (C) Cox regression analyses showing the significance and hazard ratio values of risk score and clinical characters.

### 3.5 Identification of the prognostic model-related biological processes

It is meaningful to figure out what biological processes were influenced by the prognostic risk model to make them predictive. To answer this question, functional enrichment analyses were performed. Firstly, GO enrichment was employed to analyze the DEGs between the low- and high-risk subgroups. It was observed that genes related to immune cell activation and proliferation had different expression levels (**Figure 7A**). Secondly, to further verify this observation, GSEA was utilized to find enriched pathways in the KEGG database. Results showed that 53 gene sets were significantly upregulated in the low-risk subgroup (normalized *p* < 0.05 and FDR q < 0.05) but no gene set was significantly upregulated in the high-risk subgroup (**Table S7**). Interestingly, it was observed that the most enriched biological processes in the low-risk subgroup were closely associated with immune responses (**Table S7** and **Figure 7B-G**), including the chemokine signaling pathway (NES = 2.566), Toll-like receptor signaling pathway (NES = 2.507), leukocyte transendothelial migration (NES = 2.488), T cell receptor signaling pathway (NES = 2.423), cytokine-cytokine receptor interaction (NES = 2.402), NK cell-mediated cytotoxicity (NES = 2.238), etc. These results proved that the PRGs-based prognostic risk model is related to immune responses, which indicated that the effects of pyroptosis on the immune microenvironment can influence the prognosis of SKCM.

**Figure 7:**
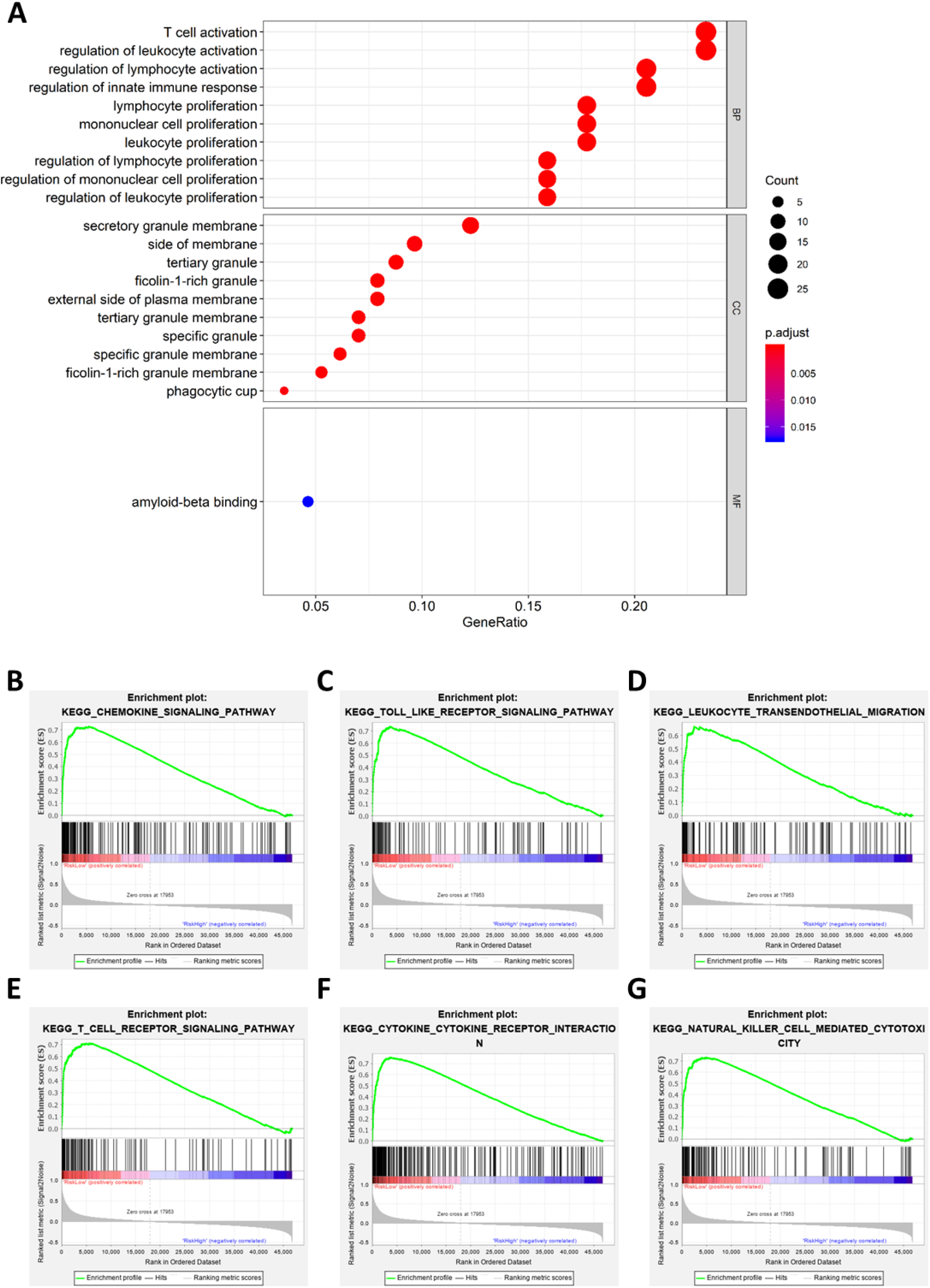
Functional enrichment analyses. **(A)** Bubble graph for GO enrichment (the bigger bubble means the more genes enriched, and the increasing depth of red means the differences were more significant). **(B-G)** Representative enrichment plots generated by GSEA reveal that the low risk was significantly associated with chemokine signaling pathway (B), Toll-like receptor signaling pathway (C), leukocyte transendothelial migration (D), T cell receptor signaling pathway (E), cytokine-cytokine receptor interaction (F), NK cell-mediated cytotoxicity (G).

### 3.6 PRGs-based prognostic risk model is related to immune responses

In order to further verify the relationship between the prognostic model and immunity, CIBERSORT was employed to evaluate the immune cell component in SKCM tissues. The proportion of 22 human immune cell subpopulations, including naive and memory B cells, plasma cells, seven T cell types, NK cells, and myeloid subsets, was assessed. Results showed that fractions of activated CD4^+^ memory T cells, γδ T cells, and M1 macrophages were significantly higher in the low-risk subgroup, whereas the high-risk subgroup had a higher fraction of M2 macrophages (**Figure 8A, Figure S5**).

**Figure 8:**
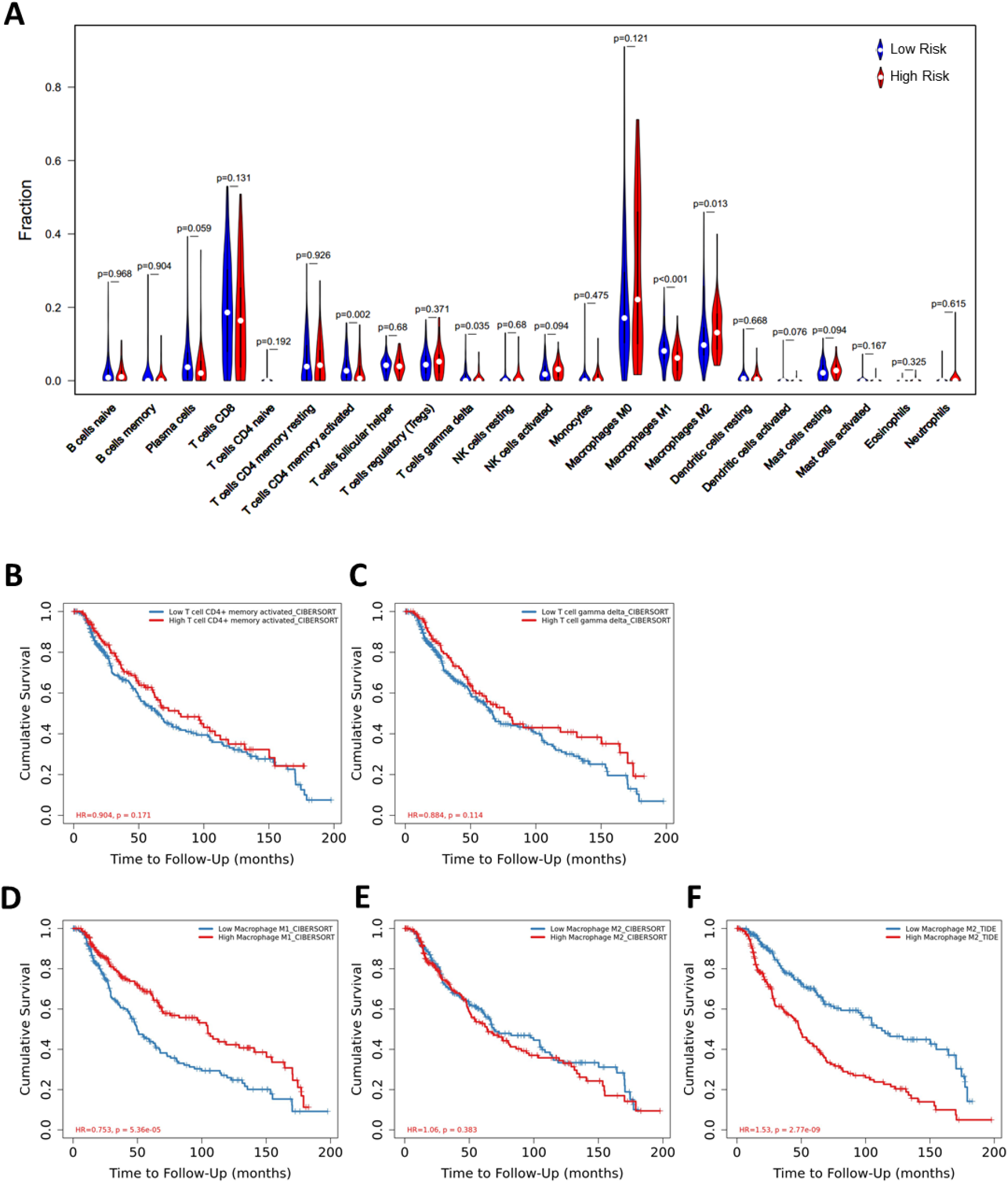
Comparison of the immune microenvironment between subgroups. **(A)** Violin plot showing the relationship between the risk score and immune fractions. The red color represents the high-risk subgroup while the blue color represents the low-risk subgroup. **(B-F)** KM curves showing the relationship between cumulative survival and immune cells in significantly different infiltrating levels, including activated CD4+ memory T cells (B), γδ T cells (C), M1 macrophages (D), and M2 macrophages (E, F). HR, hazard ratio.

Immune cells in the tumor microenvironment can influence the OS of SKCM, which may explain why the PRGs-based prognostic model can predict the prognosis. In order to prove the above hypothesis, we retrieved the relationship between immune cell infiltration and cumulative survival with Timer2.0 (**Figure 8B-F**). Interestingly, only the contents of macrophages showed significant associations with survival, where the high level of M1 macrophages or the low level of M2 macrophages indicated better survival (**Figure 8D, F**). Inflammation can be regulated by various types of tumor-associated macrophages (45). These findings suggest that, in the PRGs-based prognostic model, high-risk patients have less pro-inflammatory M1 macrophages and more anti-inflammatory M2 macrophages than low-risk patients, eventually resulting in a worse prognosis.

### 3.7 Identification of risk-related genes

Since PRGs have been shown to have prognostic significance, identifying risk-related genes would aid in further research into the function of pyroptosis in SCKM. The correlation of the prognostic risk score and the expression level of each gene was analyzed by Pearson’s correlation analysis to screen the most relevant genes. Genes with the *p* < 0.05 and the absolute value of Pearson Correlation Coefficient (|Cor|) ≥ 0.6 were considered as the strong-correlated genes (Table S8). Among them, the most relevant gene is NLRC4 which is also a component of the prognostic model. Respectively, KM survival analyses were performed for each gene with the *p* < 0.05 and |Cor| ≥ 0.7 (**Figure 9A-F**). It was observed that all the six most relevant genes were significantly associated with survival, and higher expression means longer lifespan (**Figure 9G-L**). These results imply that these genes may be involved in the pyroptosis of SKCM and function as protectors to patients.

**Figure 9:**
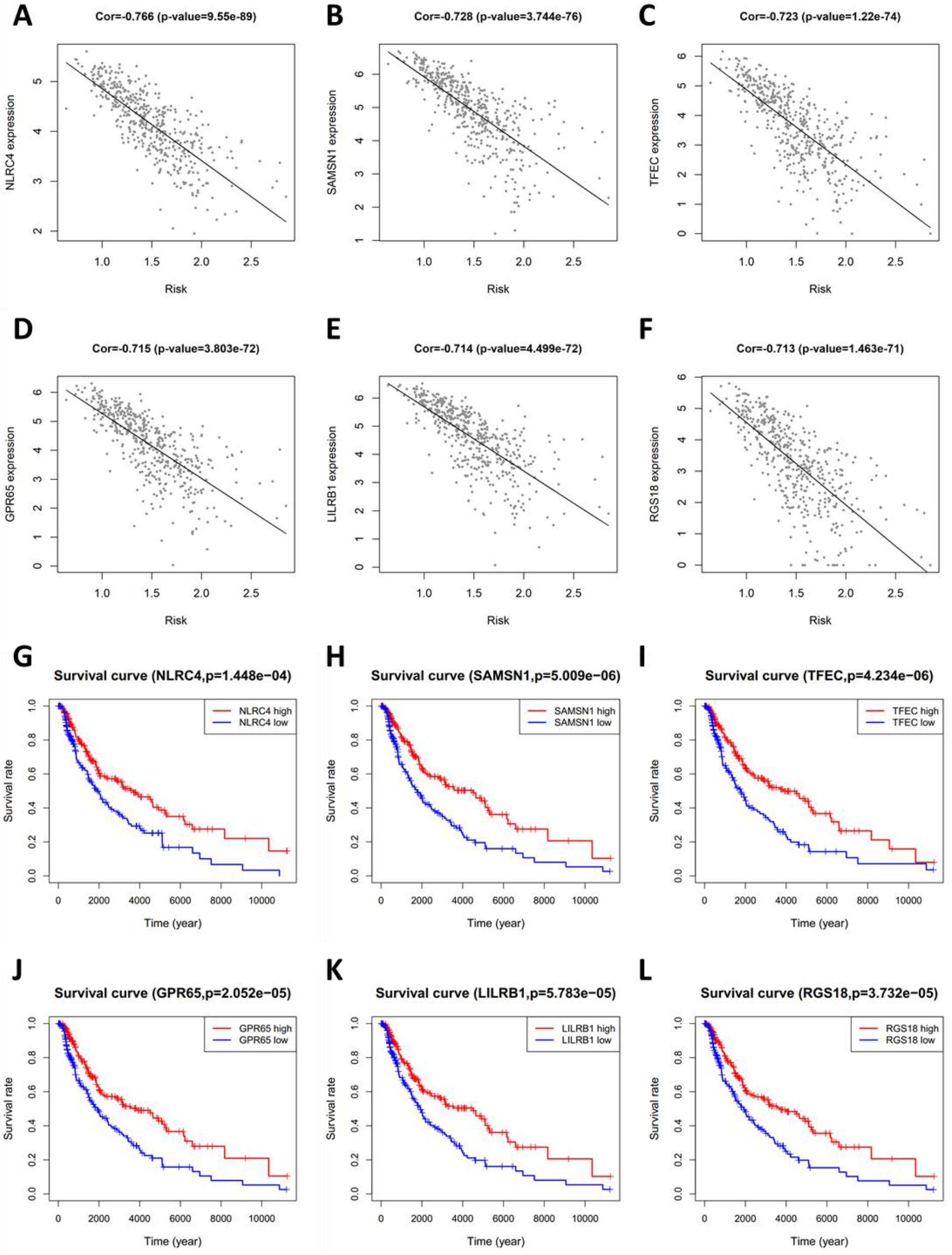
Identification of risk-related genes. **(A-F)** Representative results of correlation analysis between the risk score and each gene in SKCM. Cor: correlation coefficient. **(G-L)** KM curves showing the relationship between the six most relevant genes and OS.

## 4 Discussion

Since pyroptosis may be a double-edged sword for cancer patients, the most straightforward and concrete way to explain its importance is to develop pyroptosis-related prognostic and diagnostic models. The mRNA levels of 20 PRGs were investigated in SKCM and normal tissues in this study, and it was discovered that they were all differentially expressed. The significance of these genes related to the survival of patients was studied. Several genes that were highly expressed in SKCM and lowly expressed in normal skin tissues, but those genes were shown to be associated with a better prognosis, such as GSDMD and NLRC4, which is consistent with previous findings (46). Furthermore, diagnosis by a single gene is difficult and inaccurate. So it seems that a single PRG is unreliable for SKCM diagnosis and predicting the prognosis. This has inspired us to explore the diagnostic and prognostic value of pyroptosis by using a multi-PRG signature.

First of all, we established the SKCM-normal classifiers based on nine commonly used algorithms. Although there was little overfitting, the classifiers still had reasonable generalization ability and classification performance, especially classifiers based on the ANN, logistic regression, random forest, and SVM. Except for the decision tree, classifiers constructed from other tree-based algorithms (random forest, XGBoost, LightGBM, and Catboost) also had excellent performance. Since the PRG signature has the ability to diagnose SKCM, it is critical to collect more training samples and further tune parameters for the advancement of this SKCM diagnostic method. Clinically, because skin tissue biopsy is easier to obtain than visceral organs, PRGs-based classifiers will provide a novel and reliable method for distinguishing between malignant melanoma and benign nevus.

Secondly, we proved that PRGs expression signature has prognostic value in SKCM. To verify the hypothesis, it was found that PRGs could cluster SKCM patients, and patients in different clusters have different clinical outcomes. This suggested that the occurrence of pyroptosis in tumor tissues may be different in SKCM patients, which led to a different OS. Then we constructed a 9-gene prognostic risk model *via* LASSO Cox regression analysis, and patients in different risk subgroups had different OS, which was then validated to perform well in the external datasets.

Through the enrichment analysis of biological processes for different risk subgroups, it was found that there were significant differences in immune-related signaling pathways, which is in line with our expectations. Because the process of pyroptosis can lead to the secretion of many inflammatory cytokines, and it is also the result of inflammasome activation (6, 29). Interestingly, in addition to the representative results shown in Figure 7, we also found several signaling pathways associated with immunological rejection and autoimmune-related diseases including Type 1 diabetes. This may be due to the fact that certain patients have been treated with immune checkpoint therapy, such as ipilimumab (47, 48). While our study centered on melanoma, the importance of pyroptosis in immune checkpoint and autoimmune diseases deserves more investigation. In addition to immune checkpoint therapy, some commonly used melanoma-targeting drugs, including BRAF and MEK inhibitors, also affect the immune microenvironment through pyroptosis (49). Therefore, we hypothesize that patients will benefit from these drugs, and their curative efficacy can be monitored by PRGs-based risk score to guide the treatment.

Furthermore, pyroptosis was firstly discovered in the infectious pathogenic bacteria Shigella and Salmonella, which induced lytic cell death in macrophages by activating caspase-1 through secreted effector proteins SipB and IpaB, respectively (50, 51). As for SKCM, circulating macrophages are selectively recruited into tumors during tumor development, where they modify the tumor microenvironment. In response to numerous microenvironmental signals produced by tumor and stromal cells, macrophages change their functional phenotypes including M1 and M2. On one hand, M1 macrophages participate in the inflammatory response, pathogen clearance, and antitumor immunity. M1 macrophages have high levels of the main histocompatibility complex class I (MHC1) and class II (MHC2) molecules, which are needed for tumor-specific antigen presentation. As a result, M1 macrophages play an important role in the inflammatory response as well as antitumor immunity. On the other hand, the M2 macrophages influence the anti-inflammatory response, wound healing, and pro-tumorigenic properties. Tumor-associated macrophages (TAMs) are M2-polarized macrophages that are important modulators of the tumor microenvironment to accelerate tumor progression (52). Coincidentally, by analyzing the fraction and types of immune cells in the microenvironment, we found that M1 and M2 macrophages were different between low- and high-risk subgroups, and it does correlate with the prognosis (**Figure 8D-F**), although there are some differences between CIBERSORT and Tumor Immune Dysfunction and Exclusion (TIDE).

Finally, we analyzed genes associated with risk scores. In particular, NLRC4 was the most associated gene with the risk score, even though it was one component of the prognostic model. This suggests that NLRC4 inflammasomes may be more involved in SKCM. The risk score can be estimated using the expression level of a single NLRC4 gene since it is associated with OS in SKCM (Figure 9G). Furthermore, it was reported that Nlrc4^-/-^ mice were shown to have increased tumor development when injected subcutaneously with mouse B16F10 melanoma (53). Therefore, the impairment of NLRC4 inflammasome in melanoma cells and the function in pyroptosis are worth further study.

In conclusion, our study showed 20 PRGs differentially expressed between SKCM and normal tissues, and their association with diagnosis and prognosis. Then we showed that these genes can be used to distinguish between normal and SKCM tissues. Furthermore, the risk score derived from the prognostic model based on 9 PRGs was an independent risk factor for predicting SKCM prognosis, which was found to be related to the immune microenvironment.

## 5 Abbreviations

SKCM: skin cutaneous melanoma
PRGs: pyroptosis-related genes
DEGs: differentially expressed genes
PPI: protein-protein interaction
KNN: K-Nearest Neighbor
SVM: Support Vector Machine
ANN: Artificial Neural Network
ROC: receiver operating characteristic
AUC: area under curves
CDFs: cumulative distribution functions
PCA: principal component analysis
OS: overall survival
GSEA: gene set enrichment analysis
Cor: Pearson correlation coefficient
FDR: false discovery rate
NES: normalized enrichment score

## 6 Data availability statement

Publicly available datasets were analyzed in this study, these can be found in The Cancer Genome Atlas (https://portal.gdc.cancer.gov/) and Gene Expression Omnibus (https://www.ncbi.nlm.nih.gov/geo/).

## 8 Author Contributions

Anji Ju, Jiaze Tang, and Shuohua Chen conceived the study. Anji Ju designed the study and analyzed the data. Anji Ju wrote the manuscript, which was reviewed by Yongzhang Luo and Yan Fu. All authors read and approved the final manuscript.

