## supplementary figures and tables for "Pyroptosis-related gene signatures can robustly diagnose skin cutaneous melanoma and predict the prognosis": Supplementary materials.docx

**Supplementary Tables**

Table S1 Pyroptosis-related genes

Table S2 Major parametes

Table S3 Sample types

Table S4 Classification results

Table S5 Clinical informations

Table S6 Risk scores of each dataset

Table S7 GSEA reports

Table S8 Risk-related genes

**Supplementary Figure Legends**

**Figure S1: Differentially expressed PRGs and the correlations in expression.**

**(A)**	Heatmap of differentially expressed PRGs in TCGA-SKCM & GTEx-SKIN (* *p* < 0.05, ** *p* < 0.01, *** *p* < 0.001).

**(B)**	Bubble graph for PRGs (the bigger bubble and the increasing depth of red means higher significance).

**Figure S2: Protein levels of PRGs in SKCM**

Immunohistochemistry staining images of proteins encoded by PRGs in SKCM were retrieved from the Human Protein Atlas (www.proteinatlas.org)

**Figure S3: Tuning of k value for consensus clustering matrix.**

**(A)**	CDF curves in consensus clustering

**(B)**	Relative changes in the AUC of CDF curves.

**(C)**	Consensus clustering matrix

**Figure S4: Distribution of patients based on the risk score.**

**(A-C)**	Distribution of patients based on the risk score in training set (A), testing set (B), and validation set (C).

**(D-F)**	Distribution of survival time based on the risk score in training set (D), testing set (E), and validation set (F).

**Figure S5: Analyses of immune microenvironment**

**(A)**	Proportional histogram of the percentage of each immune cell.

**(B)**	Heatmap showing the amount of each immune cell in patients of TCGA-SKCM.

**(C)**	Heatmap showing the relationships among immune cells (the increasing depth of red means higher significance).

**Figure S6: Identification of risk-related genes.**

**(A)**	Heatmap showing the expression level of each prognostic model-related genes in patients of TCGA-SKCM.

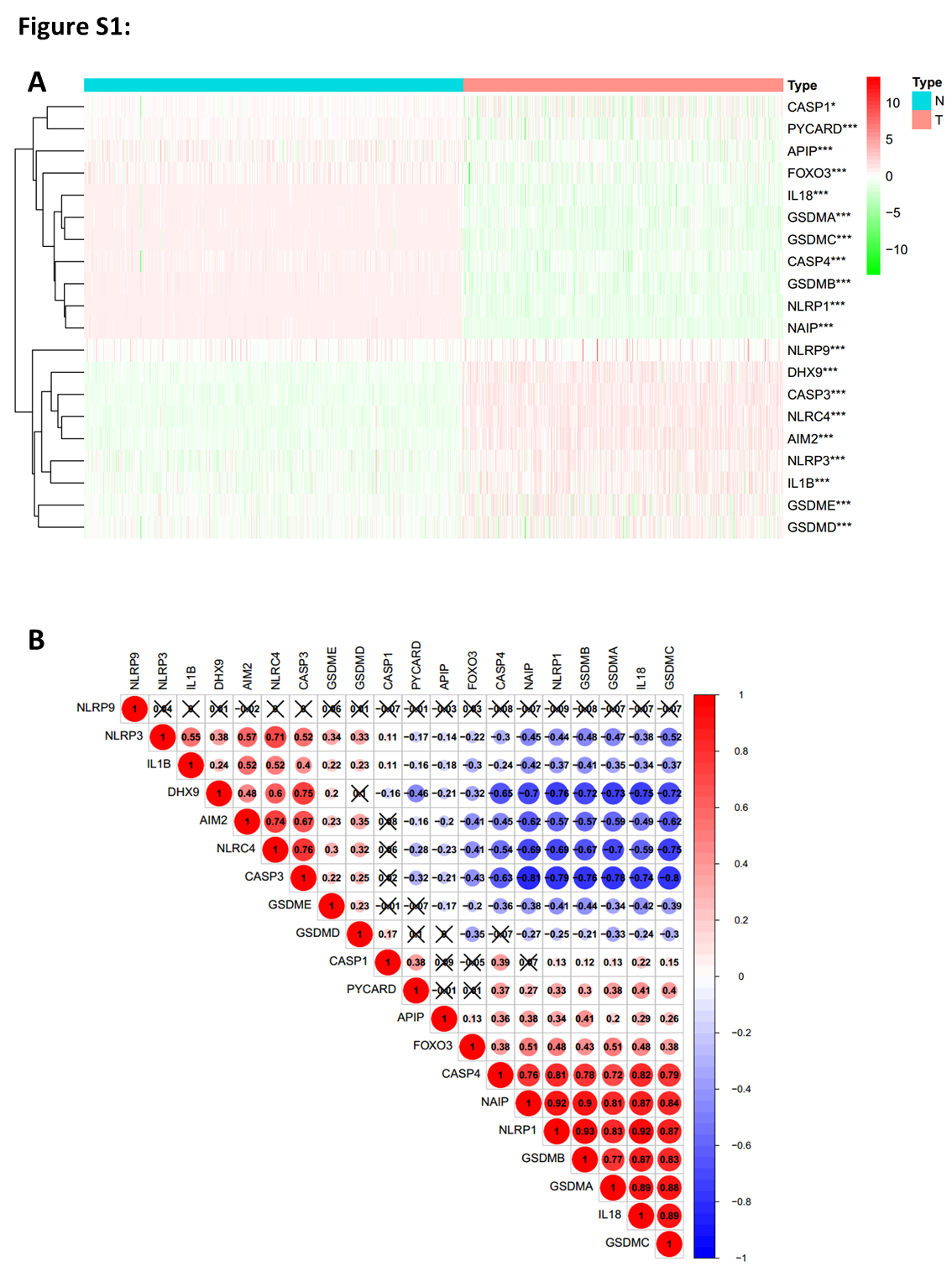

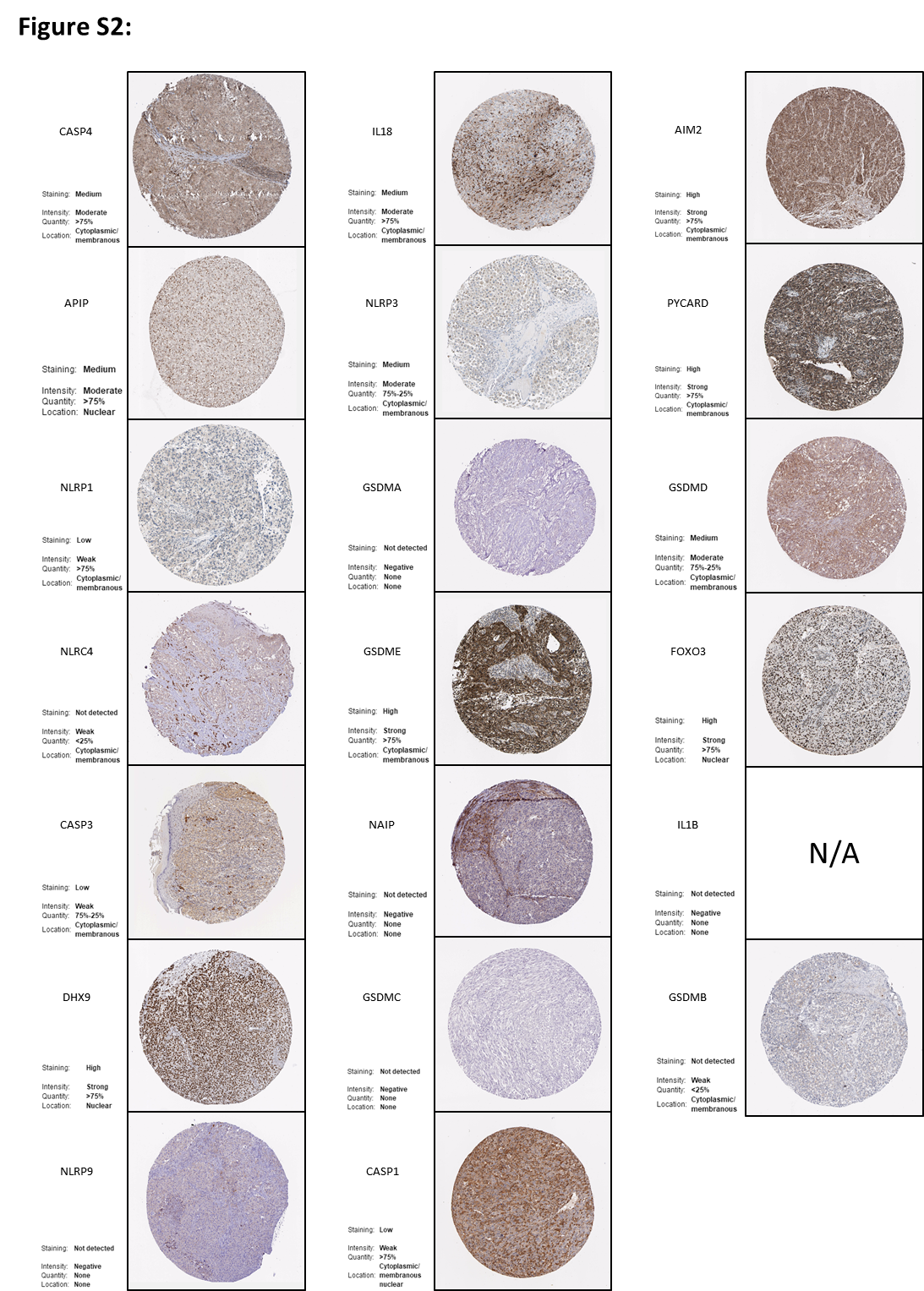

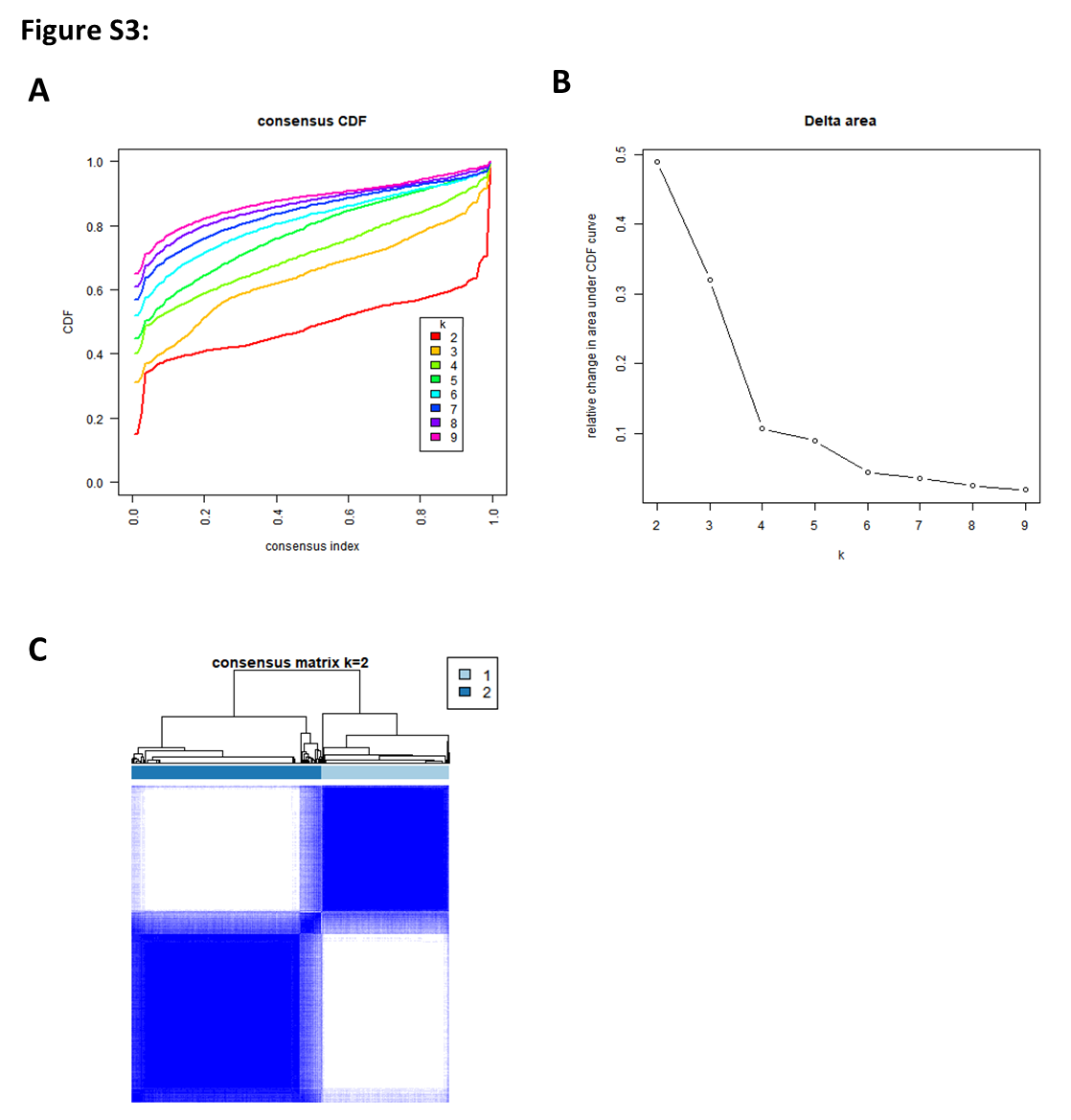

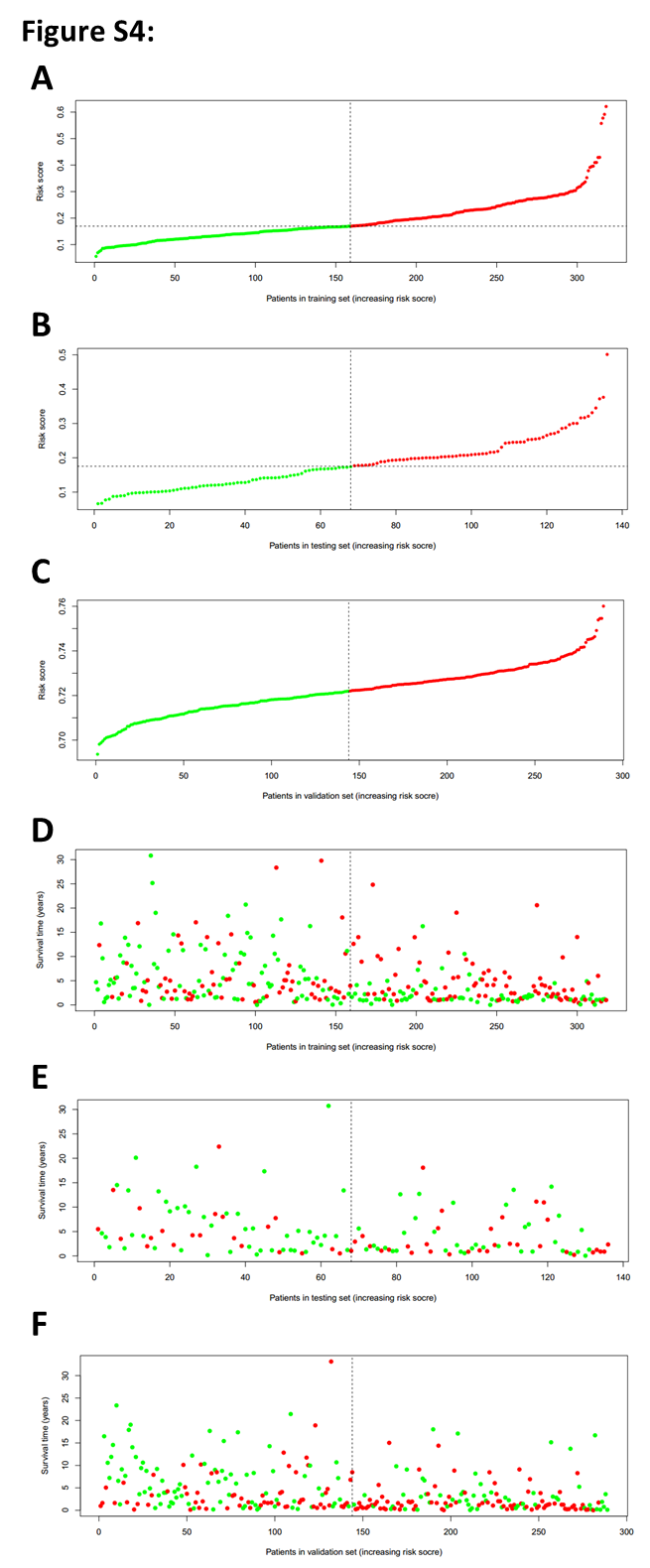

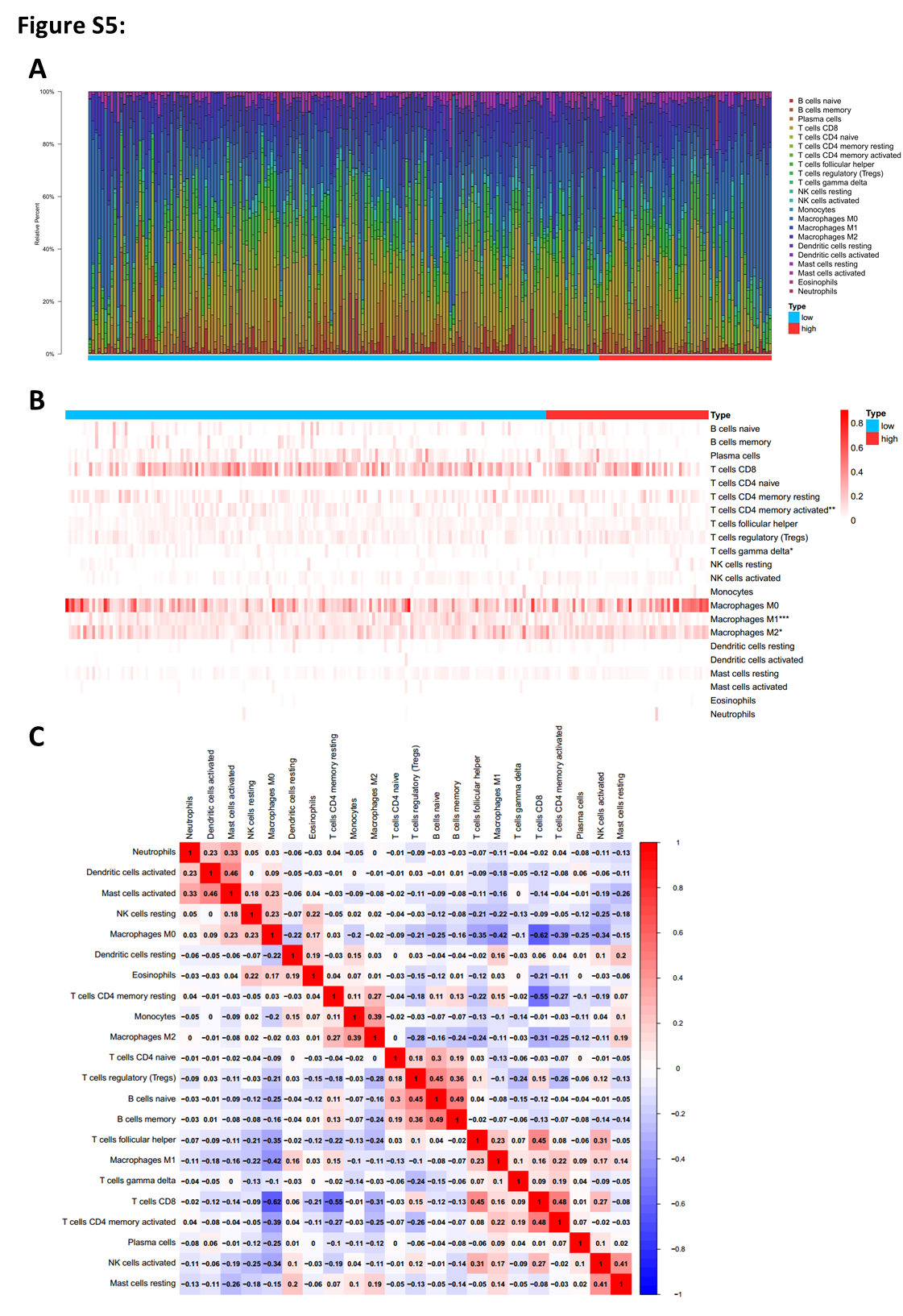
